# 6S RNA-dependent antibiotic susceptibility

**DOI:** 10.1101/2022.01.31.478597

**Authors:** Marick Esberard, Marc Hallier, Wenfeng Liu, Claire Morvan, Lionello Bossi, Nara Figueroa-Bossi, Brice Felden, Philippe Bouloc

## Abstract

Bacterial small RNAs (sRNAs) contribute to a variety of regulatory mechanisms that modulate wide ranging pathways, including metabolism, virulence, and antibiotic resistance. We investigated the involvement of sRNAs in rifampicin resistance in the opportunistic pathogen *Staphylococcus aureus*. Using a competition assay with an sRNA mutant library, we identified 6S RNA as being required for protection against low concentrations of rifampicin, an RNA polymerase (RNAP) inhibitor. This effect applied to rifabutin and fidaxomicin, two other RNAP-targeting antibiotics. 6S RNA is highly conserved in bacteria and its absence in two other major pathogens, *Salmonella enterica* and *Clostridioides difficile*, also impaired susceptibility to RNAP inhibitors. In *S. aureus*, 6S RNA is produced from an autonomous gene and accumulates in stationary phase. In contrast to what was reported in *Escherichia coli, S. aureus* 6S RNA does not appear to play a critical role in the transition from exponential to stationary phase, but affects σ^B^-regulated expression in prolonged stationary phase. Nevertheless, its protective effect against rifampicin is independent of alternative sigma factor σ^B^ activity. Our results suggest that 6S RNA helps maintain RNAP-σ^A^ integrity in *S. aureus*, which could in turn help bacteria withstand low concentrations of RNAP inhibitors.

## Introduction

*Staphylococcus aureus* is a commensal Gram-positive bacterium but also an opportunistic pathogen responsible for diseases ranging from benign (mostly cutaneous forms) to life-threatening (visceral or osteoarticular forms) infections (reviewed in (1, 2)). Due to the emergence of resistant strains, mainly methicillin-resistant (MRSA) and vancomycin-intermediate *S. aureus* (VISA), *S. aureus* has become a high priority target for the discovery of new antibiotics (3).

In standard antibiotic treatment regimens, if penicillin M and glycopeptides give unsatisfactory results, combination therapy with rifampicin may be considered, particularly in complicated prosthetic device-associated infections (4). Rifampicin, a rifamycin derivative, is an inhibitor of bacterial RNA polymerase (RNAP) (5–7). The molecule binds to the RNAP β-subunit in the DNA/RNA channel to prevent transcription by steric hindrance. This effect occurs during a narrow window, just after the synthesis of the first ribonucleotides; rifampicin is ineffective on transcripts once they are elongated (7).

Highly conserved among bacteria, the core RNA polymerase contains four essential subunits (two α, β and β’) and one accessory subunit (ω) (8–10). Among Gram-positive bacteria, RNA polymerase includes two other accessory subunits: δ and ε (ε is specific to Firmicutes). These accessory subunits may enhance transcriptional specificity and recycle RNAP. A sigma (σ) factor subunit completes the core enzyme: when present, the complex is called RNAP holoenzyme. σ factors recognize bacterial promoters and participate in adaptation to changing growth conditions (11). The σ factors are associated with specific transcriptional programs whose function and features may differ among species (11). The number of σ factors varies between species. For example, seven σ factors were described in *Escherichia coli* and *Salmonella enterica* (12), and twenty-two in the spore-forming bacterium *Clostridioides difficile* (13). In contrast, *S. aureus* possesses only four σ factors, σ^A^, σ^B^, σ^H^ and σ^S^. σ^A^ is the vegetative factor responsible for transcription of housekeeping genes (14), σ^B^ is the main alternative sigma factor contributing to stress adaptation (15–17). The last two factors are expressed only in response to specific conditions: σ^H^ is involved in the regulation of competence (18, 19) and σ^S^ in response to miscellaneous environmental stresses (20). A number of transcriptional factors participate together with σ factors in modulating bacterial transcription (21).

Small RNAs (sRNAs) are recognized as ubiquitous elements that fine-tune gene expression at transcriptional and post-transcriptional levels (22, 23). sRNAs are well studied in Gram-negative bacteria. However, in Gram-positive bacteria, including *S. aureus*, their roles in virulence, metabolism and antibiotic resistance are less understood, although there is no doubt about their involvement in these processes (24–26). The majority of characterized sRNAs interact with mRNAs. However, some sRNAs interact directly with protein complexes. This is the case for 6S RNA, one of the first described sRNAs, identified in *Escherichia coli* in 1967 (27) and sequenced in 1970 (28). In *E. coli*, 6S RNA binds preferentially to RNAP associated with the σ^70^ factor. 6S RNA accumulates during exponential growth, and reaches its maximum levels in stationary phase (29). The 6S RNA / RNAP interaction leads to inhibition of numerous *E. coli* σ^70^–dependent promoters and consequently re-orientates the transcription dependent of alternative sigma factors, allowing adaptation to many environmental conditions (reviewed in (30, 31)). Although conserved among bacteria (32), the 6S RNA role(s) and function(s) in many of them remain unknown.

We recently developed a platform to assess *S. aureus* sRNAs required for fitness based on an sRNA mutant library (33). Using this platform, we identified a rifampicin susceptibility phenotype associated with the lack of 6S RNA, pointing to a possible new mechanism of resistance against low rifampicin concentrations. We showed that this phenotype is neither restricted to rifampicin nor to *S. aureus* but extends to other RNAP inhibitors and bacterial species. Characterization of 6S RNA in *S. aureus* indicates its partial involvement in σ^B^-dependent transcription regulation at late stationary phase, rather than during transition from exponential to stationary phase. Additional experimental evidence suggests that *S. aureus* 6S RNA has a role in RNAP holoenzyme cohesion.

## Results

### Absence of 6S RNA confers increased susceptibility to rifampicin in *S. aureus* and *S. enterica*

At the beginning of this study we examined the possible involvement of sRNAs in processes underlying *S. aureus* susceptibility to antibiotics. To uncover sRNA-associated phenotypes, our laboratory has previously developed a fitness assay based on competition between sRNA-tagged mutants within a library that includes mutants of most *S. aureus bona fide* sRNAs, defined as expressed by an autonomous gene without antisense transcription (33, 34). Briefly, the fitness of individual sRNA deletion mutants growing within a collection of mutants is tested by comparing their proportion in the presence or absence of different compounds. The accumulation or reduction of individual strains are identified by monitoring the tagged sequences. This method distinguishes strains showing even subtle growth differences. Three identical libraries containing 74 putative sRNA mutants and 3 control mutants were challenged with rifampicin at a sub-lethal concentration (6 μg.L^-1^). After three days of growth, one mutant was under-represented ≃100-fold compared to the other mutants when normalized to the same libraries grown in the same medium, without rifampicin (Fig. 1). The mutant with reduced fitness due to the presence of rifampicin carried a deletion of the *ssrS* gene (referred to here as *ssrS^Sa^*), which encodes 6S RNA, an sRNA known to interact with RNAP, the rifampicin target (5–7).

**FIG 1.**
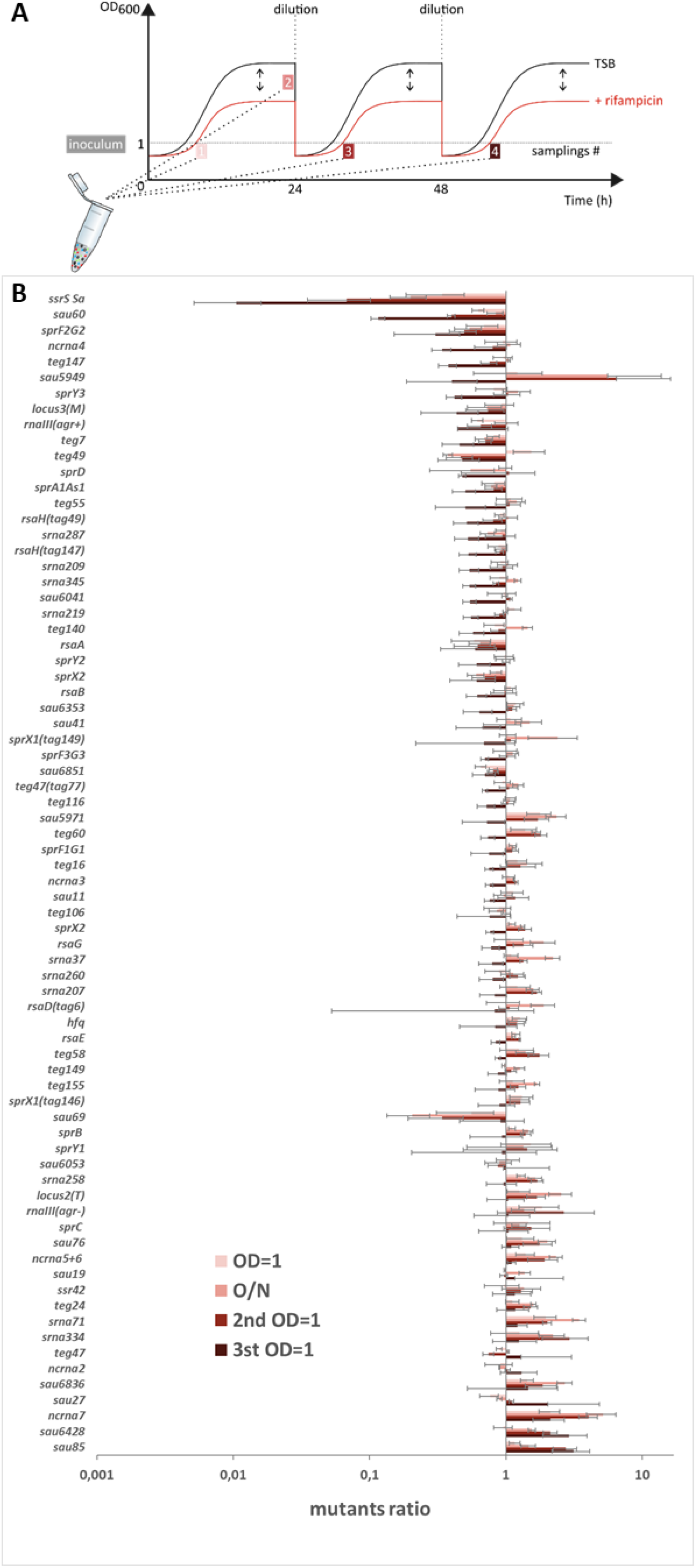
Fitness loss of the *ssrS^Sa^* mutant in the presence of sub-lethal concentration of rifampicin. (A) Scheme of the fitness experiment sampling. Three libraries were cultured during three days in Tryptic Soy Broth (TSB) +/- rifampicin. Cultures were diluted 1:1000 at 24h and 48h. After each dilution step (t= 0, 24 and 48h), samples were withdrawn for tag counting in both growth conditions, when optical density at 600 nm (OD_600_) reached the value of 1 (samplings 1, 3 and 4) and after the first overnight growth (sampling 2) as indicated. (B) Results of the competition assay between *S. aureus* sRNA mutants in presence of rifampicin 6 μg.L^-1^. y axis, mutant strain names; x axis, relative proportion of each mutant within the population grown in the presence of rifampicin normalized to the inoculum and to the corresponding sample grown in the absence of rifampicin. For each mutant, four histograms are shown; the color code corresponds to samplings indicated in Fig. 1A. Locus 2 and 3 mutants have tag insertion in loci likely not transcribed and not expected to alter the strain fitness. Error bars represent the experimental standard deviation between the three different libraries.

We asked whether *ssrS^Sa^* mutant susceptibility to rifampicin observed in the fitness experiment is detectable in monocultures. Serial dilutions of WT and *ssrS^Sa^* overnight cultures were spotted on solid medium containing low levels of rifampicin (5 μg.L^-1^); in this condition, the *ssrS^Sa^* mutant was 100-fold more sensitive to rifampicin than the parental strain (Fig. 2A). This susceptibility was reversed by insertion of an *ssrS^Sa^* copy at an ectopic chromosomal locus (Δ*ssrS^Sa^* ecto-*ssrS^Sa^*) (Fig. S1; Fig. 2A, left panel). A longer growth lag in rifampicin-containing liquid medium was also observed for the *ssrS^Sa^* mutant compared to the wild-type or complemented strains (Fig. 2C). These two tests indicated that the rifampicin susceptibility phenotype was solely due the absence of 6S RNA. This phenotype is observed within a narrow window of rifampicin concentrations, below the minimal inhibitory concentration (MIC, 12 μg.L^-1^). We conclude that 6S RNA protects *S. aureus* cells against sub-lethal concentrations of rifampicin.

**FIG 2.**
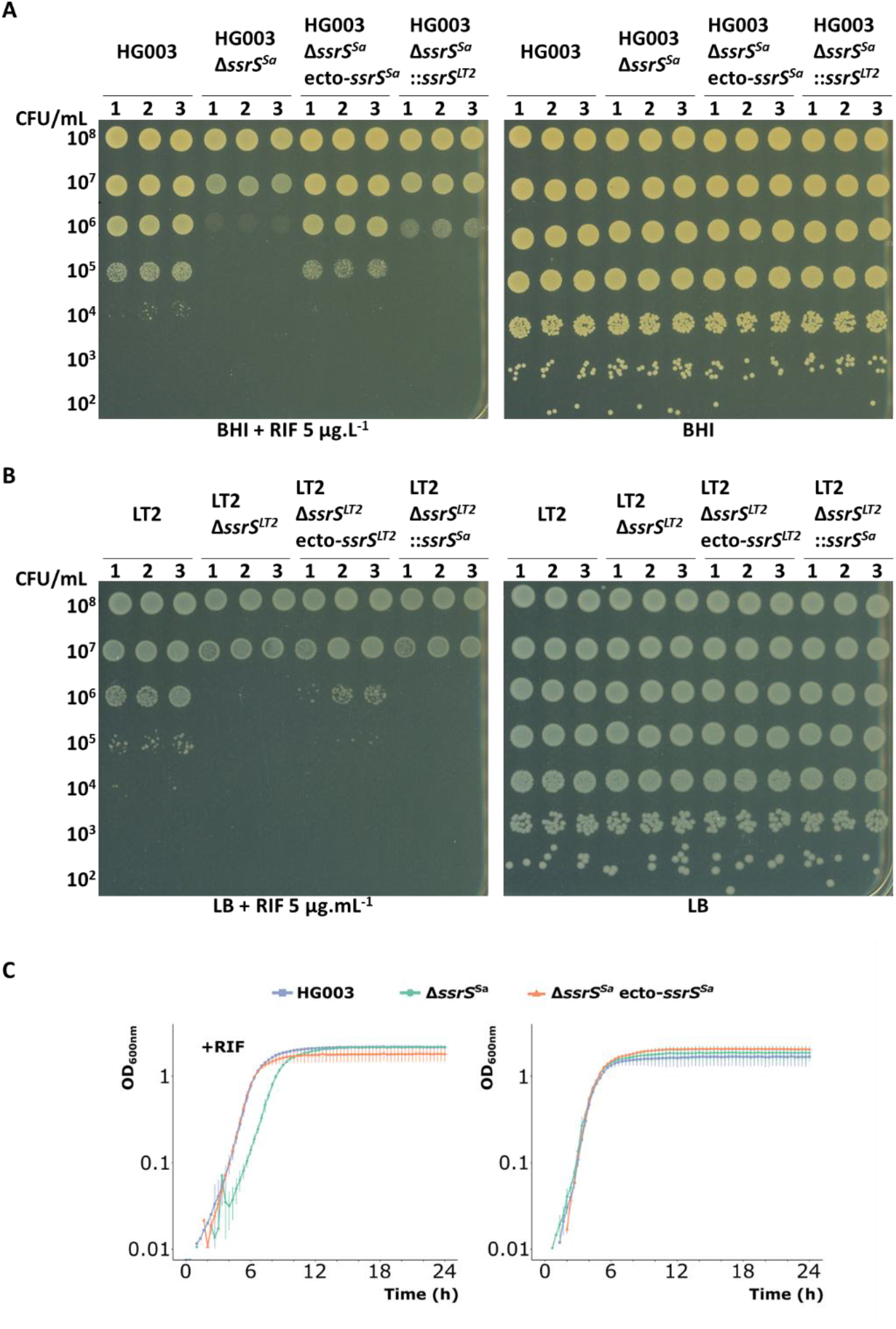
*ssrS* deletions confer a conserved rifampicin susceptibility phenotype from *S. aureus* to *S. enterica*. Three independent clones were grown overnight for each indicated strain. (A) Serial dilutions of overnight *S. aureus* cultures were spotted on Brain Heart Infusion (BHI) agar +/- 5 μg.L^-1^ rifampicin (RIF). (B) Serial dilutions of overnight *S. enterica* cultures were spotted on Lysogeny Broth (LB) agar +/- 5 μg.mL^-1^ rifampicin. (C) Growth kinetics of *S. aureus* strains in BHI +/- 5 μg.L^-1^ rifampicin. OD_600_ is an arbitrary value due to plate reader conditions, not representative of absorbance measurements of *S. aureus* in flasks. Error bars represent standard deviation of three experiments.

As 6S RNA is widely conserved in the bacterial kingdom, we examined its protective role against rifampicin in the enteric pathogen *Salmonella enterica*, a Gram-negative species. The *ssrS* gene of *S. enterica* (*ssrS^LT2^*) was deleted (Fig. S2). As in *S. aureus, ssrS^LT2^* deletion led to a rifampicin susceptibility phenotype when compared to its parental strain (Fig. 2B, left panel). However, this phenotype was only partly complemented by insertion of *ssrS^LT2^* wild-type gene at a chromosomal ectopic position (Fig. 2B, left panel).

### 6S RNA protection against rifampicin is partially interchangeable between *S. aureus* and *S. enterica*

Since *S. aureus* and *S. enterica ssrS* mutants show a similar rifampicin susceptibility phenotype, we investigated whether the 6S RNA genes would be functional in heterologous backgrounds. For this, gene swaps were performed, replacing i) the native *S. enterica* LT2 *ssrS^LT2^* gene by the *S. aureus ssrS^Sa^* homolog (*S. enterica ΔssrS^LT2^::ssrS^Sa^*; Fig S2), and ii) the native *S. aureus ssrS* gene (*ssrS^Sa^*) by the *S. enterica ssrS^LT2^* homolog (*S. aureus* Δ*ssrS^Sa^*::*ssrS^LT2^*; Fig. S1).

The *ssrS^Sa^* gene failed to compensate the *S. enterica* Δ*ssrS^LT2^* strain rifampicin susceptibility (Fig. 2B, left panel). This is possibly due reduced synthesis of staphylococcal 6S RNA in the *S. enterica* background as suggested by the results of Northern blot analysis (Fig. S3A). Interestingly, however, the reverse swap in *S. aureus ΔssrS^Sa^* partially restored growth in rifampicin (Fig. 2A, left panel). Complementation of *ssrS^Sa^* by *ssrS* from an evolutionary distant species suggests that different 6S RNAs shield against rifampicin using similar mechanisms.

### 6S RNA protects RNAP against different RNA polymerase inhibitors

The family of RNAP inhibitors comprises molecules with different mechanisms of action. We chose two RNAP inhibitors, rifabutin, fidaxomicin and a putative RNAP inhibitor, aureothricin, to test the impact of *ssrS^Sa^* on drug susceptibility.

Rifabutin, a spiro-piperidyl-rifamycin, is a rifampicin analog (35, 36). Fidaxomicin (also known as lipiarmycin (37, 38) and tiacumicin B (39)) is a narrow spectrum antibiotic (40) that inhibits transcription initiation by locking RNAP through an open-clamp state that prevents an efficient interaction with the promoter (41–44). Aureothricin is a member of the dithiolopyrrolones group and has broad-spectrum activity (45). However, the mechanism of action of this molecule remains unclear. For each drug, the appropriate sub-lethal concentrations to use were first established using the HG003 *S. aureus* strain. The *ssrS^Sa^* mutant showed a ~4 log-fold greater susceptibility to rifabutin than the HG003 strain (Fig. 3A). The *ssrS^Sa^* mutant was also negatively affected by fidaxomicin compared to the parental strain, with visibly smaller colonies (Fig. 3A). As seen with rifabutin, the susceptibility phenotype was not fully complemented by ectopic expression of *ssrS^Sa^* with either drug. Contrary to rifabutin and fidaxomicin, no aureothricin phenotype was associated with the absence of 6S RNA (Fig. 3A), possibly reflecting its poorly characterized mode of action.

**FIG 3.**
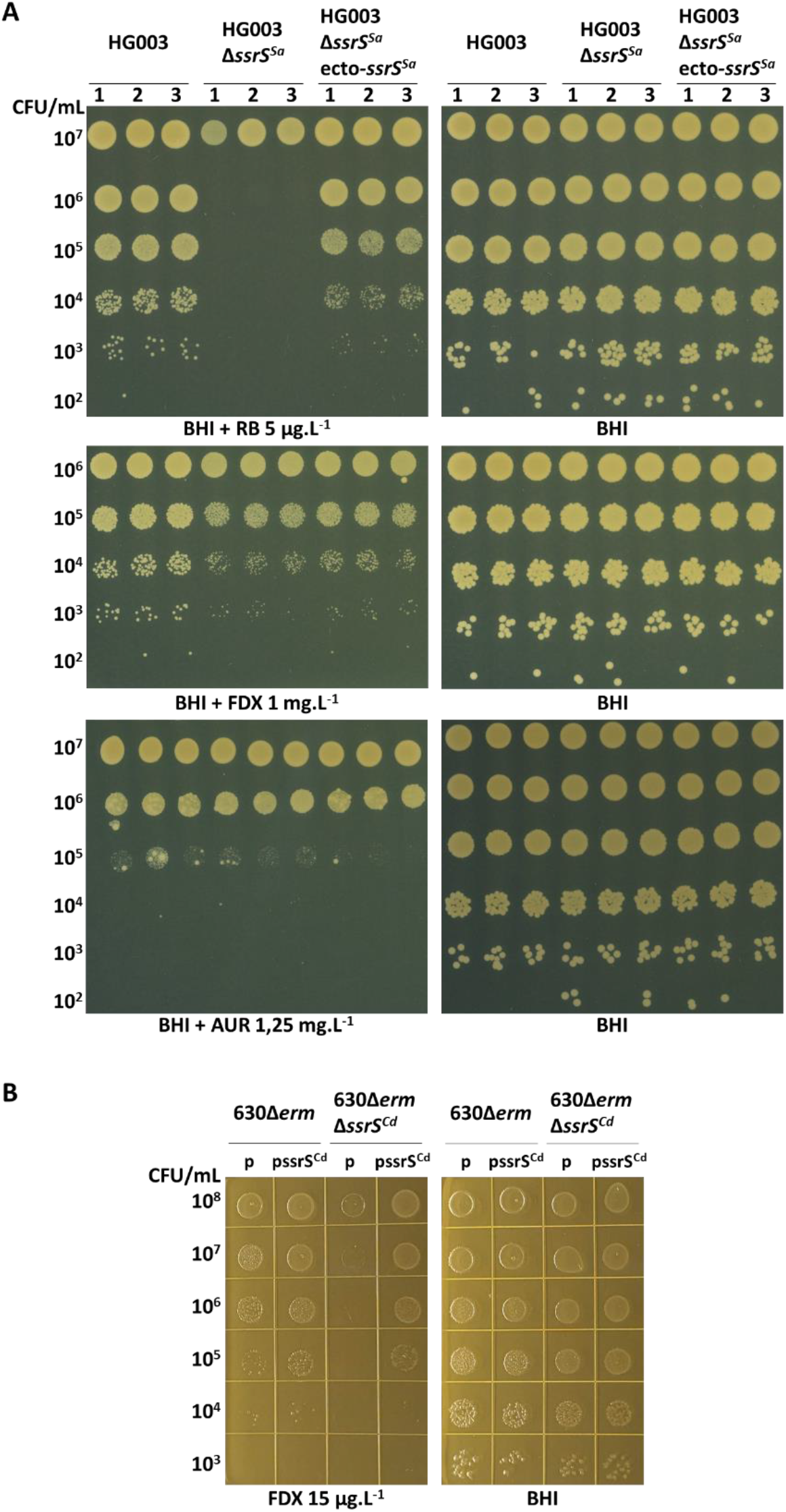
Susceptibility to RNAP inhibitors. (A) Serial dilution of *S. aureus* overnight cultures plated on solid medium containing or not antibiotics as indicated: rifabutin, RB; fidaxomicin, FDX; aureothricin, AUR; numbers (1, 2 and 3), independent clones. The antibiotic concentrations used were below the MIC. (B) Serial dilutions of *C. difficile* overnight cultures plated on solid medium containing or not fidaxomicin (FDX) as indicated. Pictures are representative of four replicates. Thiamphenicol was added in all plates (15 μg.mL^-1^) to maintain the plasmid. p, empty vector pMTL84121; pssrS^Cd^, pMTL84121-ssrS^Cd^.

Fidaxomicin is mainly active against *C. difficile*, a major human intestinal pathogen (40). We decided to test whether *ssrS* deletion also impacts RNAP inhibitor susceptibility in this species. For this, we constructed a Δ*ssrS* derivative (Δ*ssrS^Cd^*) of *C. difficile* 630Δ*erm*. *C. difficile ΔssrS^Cd^* was 1000-fold more susceptible to fidaxomicin compared to its parental strain (Fig. 3B, left panel). A plasmid carrying the *ssrS^Cd^* gene introduced in the Δ*ssrS^Cd^* strain complemented the phenotype by restoring wild-type level of susceptibility to fidaxomicin (Fig. 3B, left panel).

We conclude that *ssrS*-related susceptibility to antibiotics is a common feature of different RNAP inhibitors in evolutionary distant bacterial species. The mechanism associated with this susceptibility phenotype is likely the same for different RNAP-targeted antibiotics and different species, one possibility being 6S RNA protection of RNAP via steric occlusion.

### Growth phase-dependent expression of *ssrS* in *S. aureus*

The expression profile of 6S RNA differs according to species (30). We performed Northern experiments to evaluate 6S RNA expression in *S. aureus* (Fig. 4A). 6S RNA was strongly expressed and accumulates to 20-fold higher levels in stationary phase, as determined in *S. aureus* HG003. The expression profile of *S. aureus* 6S RNA was similar to that reported in *Salmonella* (46), *E. coli* (29) and *B. subtilis* 6S-1 RNA, which carries a second 6S RNA (47–50).

**FIG 4.**
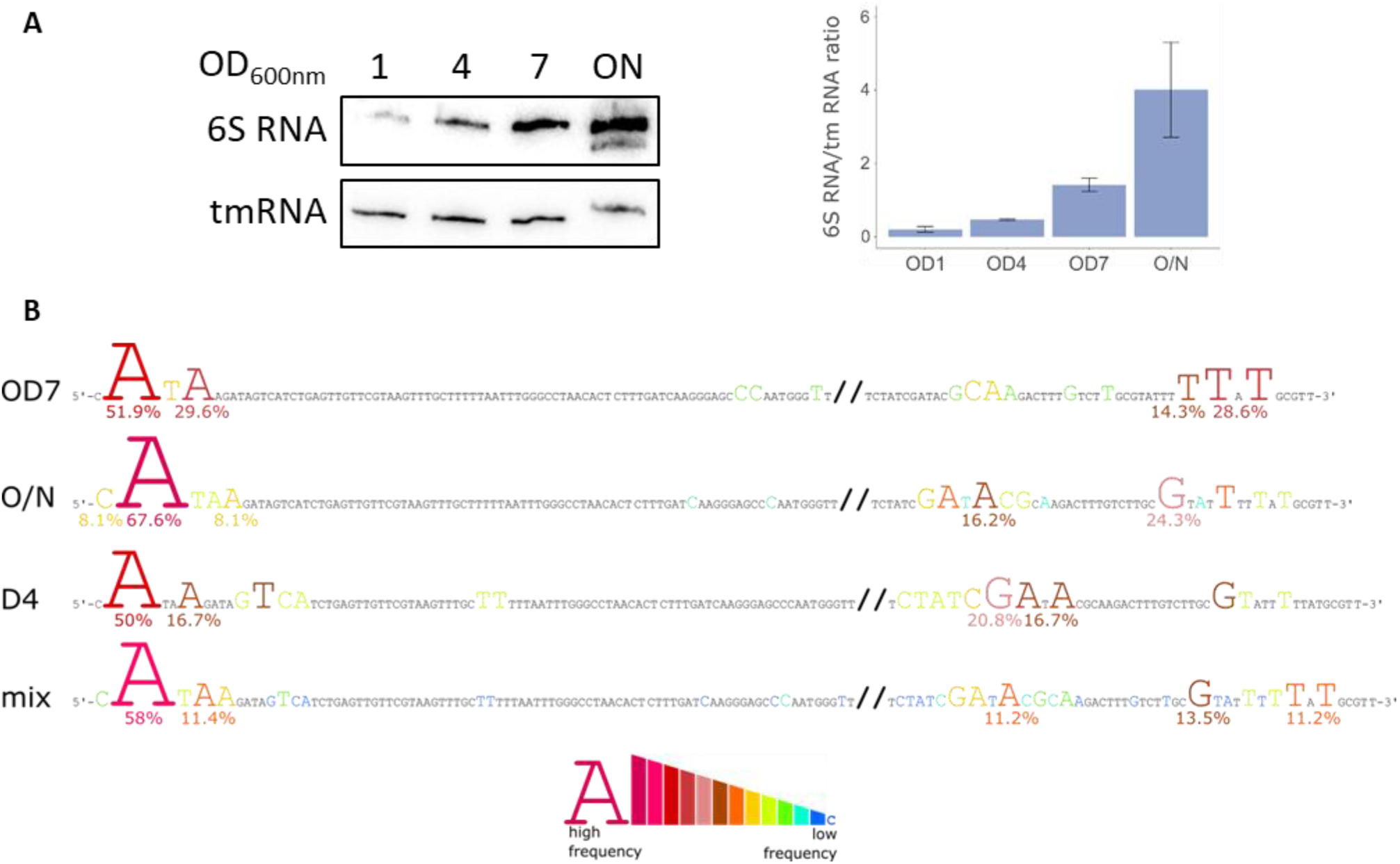
*ssrS* gene expression and 6S RNA sequence in *S. aureus*. (A) *ssrS^Sa^* expression. Cultures of HG003 grown in BHI were sampled at OD_600_=1, 4, 7 and O/N. A Northern blot probing for 6S RNA and tmRNA (for normalisation) was performed. A quantification of 6S RNA normalized to tmRNA is presented. The standard deviation is based on biological triplicates. (B) Identification of 6S RNA ends by 5’/3’-RACE mapping. Sequences were analysed separately at different time points (OD7, O/N and day4, D4) and compiled (mix). Coloured letters represent extremities found in analysed sequences. A scale of colours highlights the frequency of each 5’ or 3’-end. The highest frequencies are indicated below the corresponding nucleotides.

In overnight (O/N) samples where 6S RNA is the most abundant, a second, faster-migrating band was also observed (Fig. 4A and S3A). A second band was reported in *C. difficile* even during exponential phase (51). To determine a potential alternative 6S RNA form in *S. aureus*, also previously suggested (52), we performed 5’-3’ RACE on samples collected at different time points during growth in rich medium: OD_600_=7 (OD7, corresponding to entry into stationary phase), O/N, and on day 4 (D4). At all sampling points, the major Transcription Start Site (TSS) is the same (Fig. 4B) and in agreement with the site determined by a global TSS mapping (53). Concerning the 3’-end, the longest form ending with a T is the most abundant in OD7 samples, representing 28.6% of analyzed sequences. These data confirm that the size of the longest most abundant form is 231 nucleotides (predicted at 230 nt (52)). Samples from overnight or day 4 cultures exhibited shorter forms that may result from processing or degradation by 3’ exonucleases.

### Moderate impact of 6S RNA on the global *S. aureus* transcription profile

The role of 6S RNA in transcriptional regulation was suggested early (29) and then validated by transcriptomic analysis in *E. coli* (54–56); in RNA-seq data, 35 genes were at least 2-fold differentially expressed in a 6S RNA deficient strain compared to the parental at the onset of stationary phase. To determine whether *S. aureus* 6S RNA could play a similar role, the transcriptional profile of the Δ*ssrS^Sa^* mutant was compared with that of its parental strain by RNA-seq on samples collected at OD_600_= 7, which corresponds to the entry into stationary phase of *S. aureus* (Table 1). Transcriptome analyses were performed on biological triplicates and features with a p-value < 0.05 were retained for interpretation. Surprisingly, the transcriptional profiles of parental and Δ*ssrS^Sa^* strains were highly similar. Only three genes were more than 2-fold down-regulated in Δ*ssrS^Sa^* (FC < 0.5; Table 1 top panel). They encode a hypothetical epoxyqueuosine reductase (QueH/SAOUHSC_02911), a hemin transporter (HrtA/SAOUHSC_02640 (its co-functional partner HrtB/SAOUHSC_02641 is also down-regulated), and a 30 amino-acid peptide (SAOUHSC_01817) of unknown function. Other genes related to transporters, cell wall metabolism and redox state were also significantly reduced but with a lower-fold change. All of these genes are regulated by σ^A^, except *bstA*, a σ^B^ DNA-damage-induced gene coding a putative DinB superfamily protein. Taken together, these results suggest that 6S RNA does not redirect transcription during the stationary phase transition in a sigma-dependent manner.

**Table 1.** Transcriptomic analysis of Δ*ssrS^Sa^* vs. HG003 in late exponential phase. Fold change (FC) represents the gene expression ratio between Δ*ssrS^Sa^* and its parental strain at OD_600_ of 7. Genes with FC < 0.66 (top panel) or > 1.5 (down panel) are presented. Transcription unit (TU) are indicated by letters. Italic characters, hypothetical proteins; P-adj, adjusted p-value.

| Locus tag / gene name | FC | P-adj | Function / relevant data | Classification | Regulation | TU |
| --- | --- | --- | --- | --- | --- | --- |
| SAOUHSC_02911 | 0.48 | 2.2E-28 | *Epoxyqueuosine reductase* | tRNA modification | $\sigma^A$ | |
| SAOUHSC_02640 / *hrtA* | 0.48 | 4.2E-10 | Hemin efflux ATP-binding protein HrtA | transport and binding | $\sigma^A$, HssR | b |
| SAOUHSC_01817 | 0.49 | 5.9E-10 | *integral component of membrane* | | $\sigma^A$ | c |
| SAOUHSC_01736 | 0.52 | 1.0E-08 | *Unknown.*<br>Downstream *ssrS* (34) | | $\sigma^A$ | a |
| SAOUHSC_02641 / *hrtB* | 0.53 | 3.7E-08 | Hemin efflux system permease protein HrtB | Transport and binding | $\sigma^A$, HssR | b |
| SAOUHSC_01818 / *ald2* | 0.55 | 8.0E-07 | Alanine dehydrogenase | Energy metabolism | $\sigma^A$, CcpA | c |
| SAOUHSC_00874 | 0.56 | 3.5E-10 | *Thioredoxin-like protein* | Hypothetical protein | $\sigma^A$ | |
| SAOUHSC_02297 | 0.57 | 3.7E-14 | S1 RNA-binding domain-containing protein | Protein synthesis | $\sigma^A$ | d |
| SAOUHSC_02590 | 0.60 | 3.7E-09 | *Amino acid permease* | Transport and binding | $\sigma^A$, CcpA, CodY | |
| SAOUHSC_02296 | 0.62 | 1.7E-08 | *SprT-like protein* | | $\sigma^A$ | d |
| SAOUHSC_00561 / *vraX* | 0.64 | 4.2E-10 | *VraX* | | $\sigma^A$ | |
| SAOUHSC_00704 | 0.64 | 2.0E-08 | *ABC-2 transporter* | | $\sigma^A$ | |
| SAOUHSC_03028 / *bstA* | 0.64 | 1.2E-03 | *DinB-like protein* | | $\sigma^B$ | |
| SAOUHSC_01735 / *tcdA* | 0.65 | 1.9E-06 | *ThiF domain-containing protein* | Cofactors biosynthesis | $\sigma^A$ | a |
| SAOUHSC_02656 | 0.65 | 4.0E-10 | *Cytochrome c oxidase-like protein* | | $\sigma^A$ | |
| SAOUHSC_00157 / *murQ* | 1.51 | 1.6E-03 | N-acetylmuramic acid-6-phosphate etherase | Cell envelope | $\sigma^A$, MurR, CcpA | e |
| SAOUHSC_00156 / *mupG* | 1.52 | 1.2E-03 | *6-phospho-N-acetylmuramidase* | Cell envelope | $\sigma^A$, MurR, CcpA | e |
| SAOUHSC_01121 / *hla* | 1.56 | 1.6E-05 | Alpha-hemolysin | Virulence / toxin | $\sigma^A$, SaeR, CcpA, RNAIII | |
| SAOUHSC_02169 / *chp* | 1.60 | 1.8E-04 | Chemotaxis-inhibiting protein CHIPS | Virulence | $\sigma^A$ | |
| SAOUHSC_00961 / *comK1* | 1.85 | 4.3E-08 | *Competence protein* | | $\sigma^A$, CodY | |

Transcriptome results, as performed in the early period of stationary phase, did not provide evidence linking σ^B^ transcriptional activity to the presence of 6S RNA. We used a reporter fusion strategy to pursue this question: the gene encoding the fluorescent protein mAmetrine was placed under the transcriptional control of the σ^B^-regulated *SAOUHSC_00624* promoter (57) (pPsigB-mAmetrine; Fig. 5A). No significant difference of fluorescence was observed between *ssrS^Sa^* and parental strains when measured for the first 18h of growth in rich liquid medium. Thus, in keeping with transcriptomic findings, we conclude that 6S RNA does not appear to redirect transcription during transition from exponential to stationary phase in *S. aureus*, which differs from what was reported in *E. coli* (56–58). Interestingly, however, after 18h of culture, mAmetrine expression continues to increase in the parental strain, compared to markedly lower expression in Δ*ssrS^Sa^*. This result, suggesting that 6S RNA could be important for efficient σ^B^-dependent gene expression during starvation, remains to be investigated.

**FIG 5.**
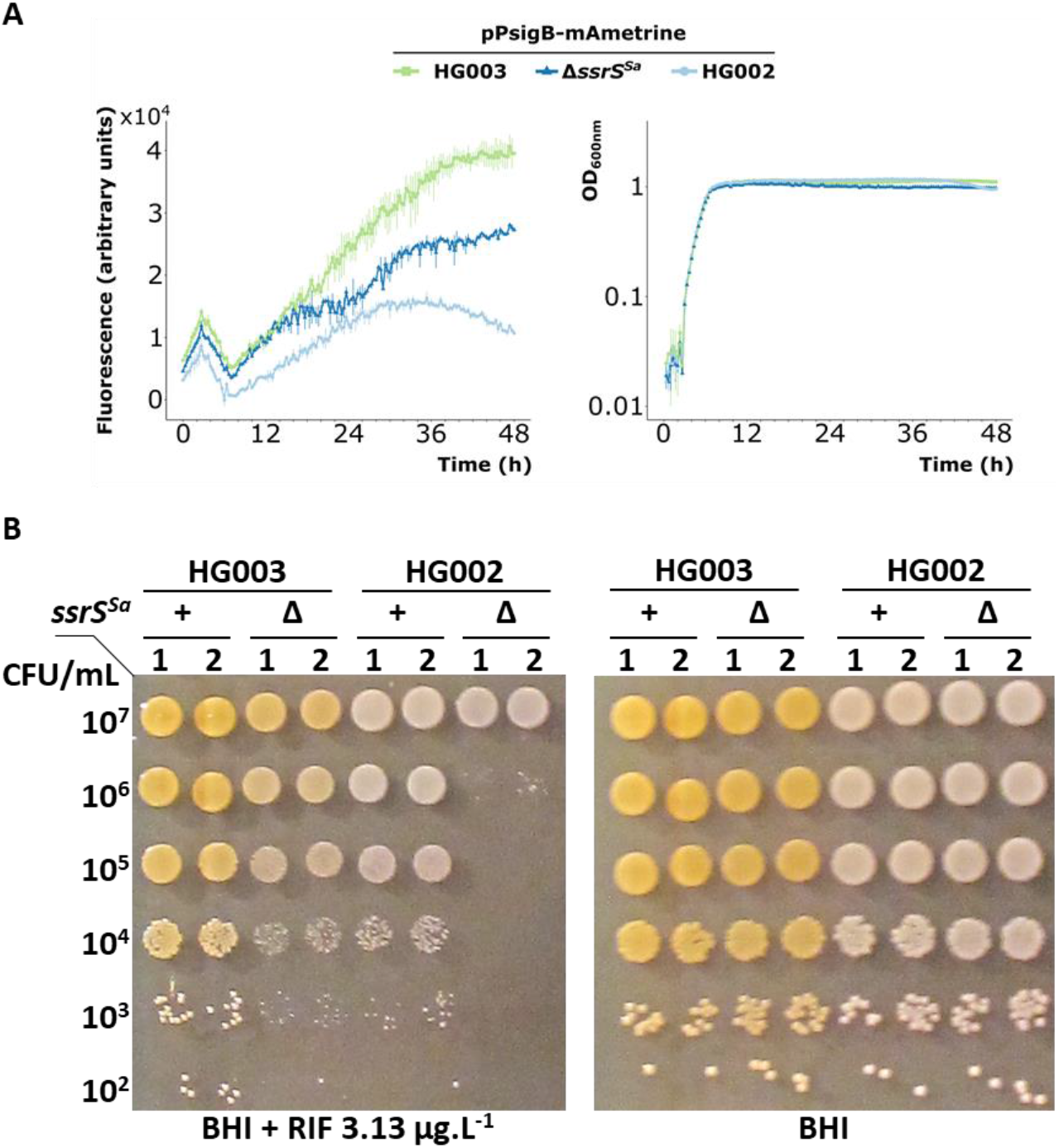
6S RNA and σ^B^ interplay in late stationary phase and in rifampicin response in *S. aureus*. (A) Fluorescence and OD_600_ were monitored simultaneously in three strains (HG003 [parental], HG003 Δ*ssrS^Sa^* and HG002) expressing a fluorescent protein (mAmetrine) under the control of the σ^B^ promoter of *SAOUHSC_00624* from a plasmid (pPsigB-mAmetrine). HG002, *rsbU*^-^ strain equivalent to σ^B-^ strain, is a negative control. Error bars represent standard deviation of biological triplicates. (B) Spot test comparing HG003 and HG002 (parental and Δ*ssrS^Sa^* strains, respectively) with sub-lethal concentration of rifampicin (3.13 μg.L^-1^). Arbitrary values are shown as OD_600_. Experiment was done with independent duplicates.

Since the absence of 6S RNA in *S. aureus* leads to an increased susceptibility to rifampicin, we questioned if this phenotype was related to σ^B^ regulation. The HG002 strain is a HG003 isogenic strain containing an 11 bp-deletion in the *rsbU* gene encoding a σ^B^ activator (59). Consequently, HG002 is deficient in σ^B^ activity, illustrated by the absence of yellow pigmentation (60). We first noticed a greater susceptibility to rifampicin of HG002 compared to HG003 (Fig. 5). This observation indicates that lower σ^B^ activity *per se* confers increased rifampicin susceptibility, as described in *C. difficile sigB* mutant (61). This effect is probably due to a stress adaptation deficiency related to the absence of σ^B^ regulation. To determine the effect of 6S RNA in this genetic context, the *ssrS^Sa^* deletion was introduced in HG002. The resulting strain (HG002 Δ*ssrS^Sa^*) was considerably more susceptible to rifampicin than the parental strain HG002 (Fig. 5B). This observation indicates that the absence of 6S RNA leads to increased rifampicin susceptibility through a pathway that is independent of σ^B^ activity.

### 6S RNA plays a role in RNAP stability in *S. aureus*

6S RNA binds to RNAP-σ^70^ in *E. coli* (29, 47). As RNAP holoenzyme is a protein complex with accessory subunits (especially σ factors), an element binding to this complex could directly influence its stability or composition. We first performed an electrophoretic mobility shift assay (EMSA) in *S. aureus* with radiolabeled 6S RNA (P^32^-6S RNA), purified sigma factors (σ^A^-His and σ^B^-His) and RNAP (with His-tagged RpoC) to assess the interaction with 6S RNA (Fig. 6A). A P^32^-labelled unrelated sRNA, SprB (62), was used as a control. No interactions were detected with SprB. In contrast, our results showed interaction between 6S RNA and RNAP coupled to the vegetative sigma factor, σ^A^, to a lesser extent with RNAP-σ^B^.

**FIG 6.**
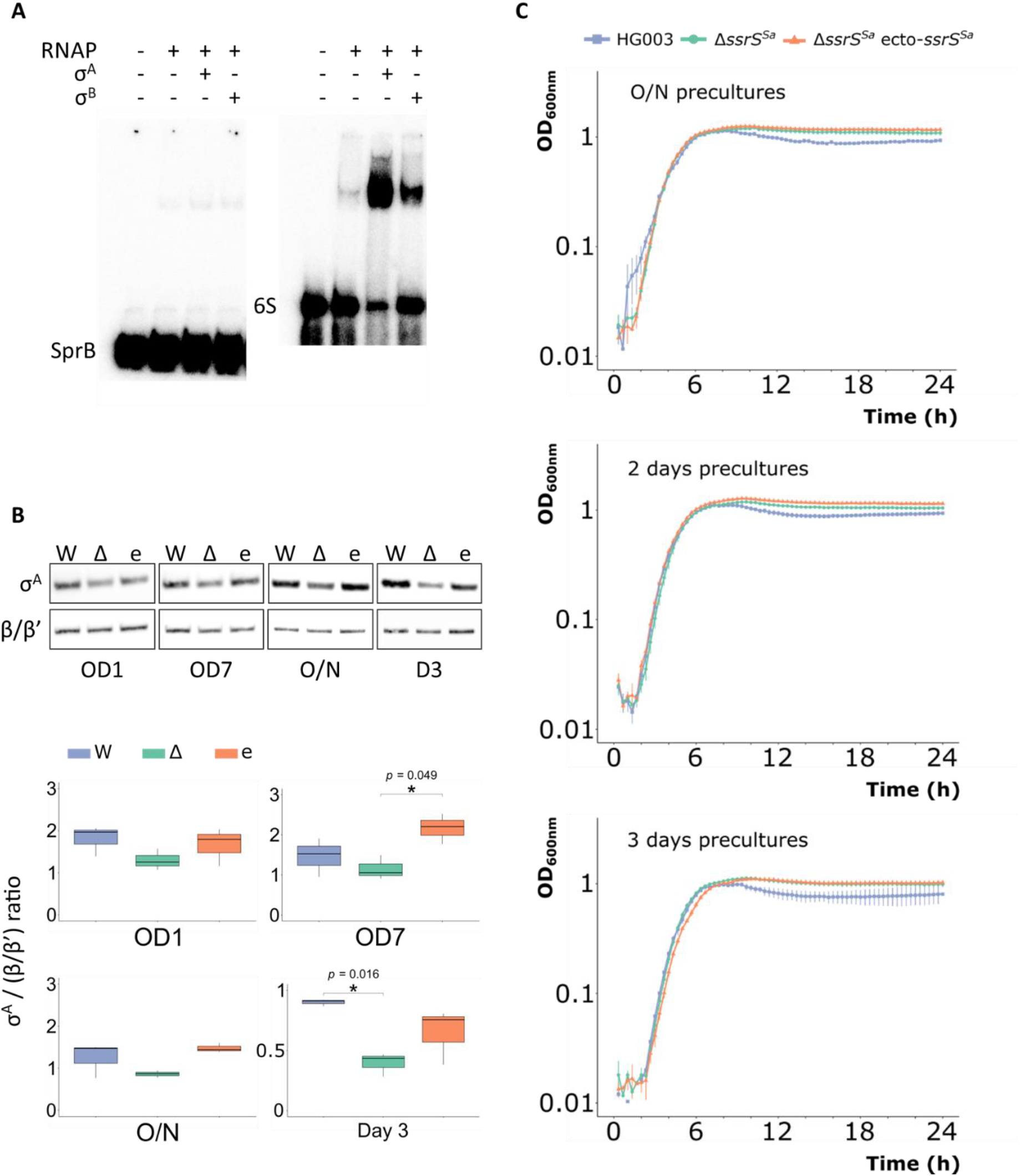
6S RNA and RNAP holoenzyme interactions in *S. aureus*. (A) EMSA with P^32^-6S RNA (6S RNA), RNAP, σ^A^ and σ^B^. All the proteins are His-tagged purified. P^32^-SprB (SprB) is a control RNA. (B) Immunodetection of σ^A^ performed by Western-blot from samplings at OD_600_=1 and 7, O/N and day 3 (D3). Quantification of σ^A^ is relative to the amount of RNAP β/β’ subunits. Experience realized in biological triplicates. W: wild-type (HG003); Δ: *ssrS^Sa^* mutant; e: Δ*ssrS^Sa^* ecto-*ssrS^Sa^*. (C) Growth curves of HG003, *ssrS^Sa^* mutant (Δ*ssrS^Sa^*) and complemented (Δ*ssrS^Sa^* ecto-*ssrS^Sa^*) strains in BHI. Strains were cultured in independent triplicates from O/N, two days or three days precultures. Error bars represent standard deviation.

We questioned if the absence of 6S RNA could alter RNAP holoenzyme composition. The amounts of σ^A^ and β/β’ subunits were evaluated in *ssrS^Sa^* and parental strain cultures at different time points. Western blots were performed with antibodies raised against σ^A^ and RNAP (Fig. 6B). Interestingly, σ^A^ pools were lower in the *ssrS^Sa^* mutant compared to the parental strain and complemented strains in all tested conditions. At day 3 (D3), σ^A^ pools were decreased by nearly 2-fold in the *ssrS^Sa^* mutant compared to the parental strain. These results suggest that 6S RNA plays a role in RNAP holoenzyme stability and could act as a protective belt for RNAP-σ^A^. We hypothesized that a reduced amount of σ^A^ could modify strain outgrowth. Growth from HG003 (parental strain), HG003 Δ*ssrS^Sa^* and its complemented mutant HG003 Δ*ssrS^Sa^* ecto-*ssrS^Sa^* from pre-cultures that had grown O/N, 2 days and 3 days were compared (Fig. 6C). Surprisingly, no growth difference was observed between the three strains in BHI, regardless of the pre-culture age. Despite the significant effect on σ^A^ levels, 6S RNA is not an essential factor for *S. aureus* growth in rich medium.

## Discussion

Here we demonstrated that the absence of 6S RNA in *S. aureus* leads to a fitness loss in the presence of low rifampicin concentrations. This marked phenotype was associated with only one sRNA gene (*ssrS^Sa^*) out of 77 tested mutants in a competition experiment. This phenotype is conserved from Gram-positive to Gram-negative bacteria suggesting a common protective effect.

In *S. aureus*, the rifampicin phenotype was fully restored by ectopic gene complementation, indicating that it was solely due to the absence of 6S RNA. In *S. enterica* however, similarly done complementation of the *ssrS^LT2^* deletion was only partial, while the native and ectopic copies had similar expression levels (Fig. S3A). In *E. coli,* the *ssrS* and *ygfA* genes are in an operon (48, 63) and mature 6S RNA results from a 5’- and 3’-transcript ends processing (64, 65); similar organisation and regulation is expected in *S. enterica.* Two hypotheses may explain the incomplete complementation of *ΔssrS^LT2^* rifampicin phenotype: (i) The ectopic *ssrS^LT2^* copy is subjected to a slightly different processing pathway (not detected in the gel in Fig. S3A) and (ii) Δ*ssrS^LT2^* could exert a polar effect on *ygfA* expression, however, no growth defect has been observed for the mutant so far.

Despite weak similarity between *S. aureus ssrS^Sa^* and the cognate *S. enterica ssrS^LT2^* gene, we observed partial complementation of the rifampicin phenotype in *S. aureus* Δ*ssrS^Sa^* by the Δ*ssrS^LT2^* allele. The lack of the reverse complementation (*ssrS^Sa^* into *S. enterica* Δ*ssrS^LT2^*) might be ascribable to the lower expression of *ssrS^Sa^* in *S. enterica* (Fig. S3A) and/or to any of the hypothesis raised above for the *S. enterica* Δ*ssrS^LT2^* ecto-*ssrS^LT2^* phenotype.

Susceptibility of the *ssrS* mutants was not observed for all the compounds tested. Differences in the mechanisms of action, binding sites and drug entry efficiencies could explain this observation (42, 66). The Δ*ssrS^Sa^* mutant showed increased susceptibility of *S. aureus* to rifampicin, rifabutin, and fidaxomicin. Similarly, cognate Δ*ssrS S. enterica* and *C. difficile* mutants showed marked sensitivities to rifampicin and fidaxomicin, respectively. These drugs bind RNAP close to sites interacting with DNA, suggesting that 6S RNA interaction with the enzyme may, at least partially, prevent antibiotic access to their sites. Based on this reasoning, our results suggest that aureothricin, for which toxicity was unaffected by *ssrS^Sa^* deletion, does not bind RNAP at the interface with DNA.

Our findings suggest differences in the regulatory roles of 6S RNA in *S. aureus* compared to those reported in *E. coli* (54). In the latter species, 6S RNA interaction with the RNAP holoenzyme is proposed to coordinate transcriptional regulation with growth (29). Accordingly, numerous transcriptome analyses performed under different conditions, indicated that in *E. coli*, many of 6S RNA-regulated genes were related to translational/transcriptional machinery or amino acid metabolism (29, 54–56, 67–70). In contrast, our *S. aureus* transcriptomic analysis and promoter assay reveal no obvious 6S RNA-related differential expression during transition to stationary phase. Two major features of *S. aureus* could explain this phenomenon: (i) lower diversity of sigma factors in *S. aureus* with only four σ factors, among which σ^A^ and σ^B^ control the majority of transcribed genes. Knowing that the main alternative sigma factor σ^B^ is involved in stress response and not only in stationary phase adaptation, σ^B^-promoters could be less sensitive to 6S RNA during the transition phase; (ii) the compensatory effect of a co-regulator. The levels of the alarmone ppGpp increases in *E. coli ssrS* mutants, and might compensate the lack of 6S RNA (55, 56, 58). This possibility provides an attractive explanation in *S. aureus*, as ppGpp is synthesized in response to nutrient starvation, and drives growth adaptation (71). Further experiments are needed to explore this pathway in *S. aureus*.

Our promoter assay (Fig. 5A) suggests a 6S RNA-dependent expression of σB-promoters in late stationary phase, after 18h of culture. Among its known roles in *E. coli*, 6S RNA also influences transcription during long-term starvation (54, 67). Whether *S. aureus* 6S RNA interacts with σ^B^ for alternative promoter expression during late stationary phase remains unclear. In particular, in the absence of a functional σ^B^, the absence of 6S RNA still generates rifampicin inhibition indicating that this phenotype was not due to a lack of reprograming transcription from σ^A^ to σ^B^. The relationship between 6S RNA and σ^B^ remains to be characterized.

We showed that *S. aureus* 6S RNA interacts directly with RNAP-σ^A^, raising the question of whether this could directly affect the holoenzyme stability. Of note, the *Δsau60* mutant exhibits a moderate reduction in fitness in the presence of rifampicin (Fig. 1B). Δ*sau60* corresponds to a deletion within the intergenic sequence upstream of *rpoB* encoding the β subunit of RNA polymerase; this deletion may alter the ratio of RNAP subunits and possibly the RNAP stability, leading to a rifampicin phenotype. However, this attenuated phenotype was not detected by a spot test. In *E. coli*, the majority of 6S RNA is coupled to RNAP-σ^70^ (29). In *Streptococcus pneumoniae,* 6S RNA bound to RNAP was recently proposed to be a stockpile for inactive RNAP (72). In *S. aureus*, the absence of 6S RNA leads to a reduced amount of σ^A^ in prolonged stationary phase cultures. A similar effect was observed with σ^70^ in *E. coli* (29) and in soluble sigma fraction of *Synechocytis* sp. (73). In *S. aureus*, σ^A^ is unstable (74); our results indicate that it is probably stabilized by RNAP core enzyme and 6S RNA. Knowing that σ^A^ is the vegetative sigma factor in *S. aureus*, decreased levels in the Δ*ssrS^Sa^* strain could have a negative impact on growth, and particularly on outgrowth recovery. As comparison, in *B. subtilis*, which expresses two different 6S RNAs, outgrowth is delayed in cells lacking 6S-1 RNA (75), whereas no extended lag phase was noticed in *E. coli 6S* RNA deficient cells (29). Similar to *E. coli* (29), no lag linked to *ssrS* was observed during *S. aureus* outgrowth from stationary phase, suggesting that reduced σ^A^ pools in the *ssrS^Sa^* mutant are enough to manage growth restart and that 6S RNA is not essential for growth in rich medium.

RNAP inhibitors remain in use in combination therapies against difficult-to-treat infections (4). Antibiotic concentrations below the MIC are encountered by bacteria in many environmental conditions including hosts undergoing antimicrobial treatments (76). We demonstrated that 6S RNA provides protection against low concentrations of RNAP inhibitors. 6S RNA is highly conserved, and the effects of *ssrS* deletion on RNAP inhibitor susceptibility were observed in unrelated pathogens. 6S RNA may significantly enhanced fitness to RNAP inhibitors under these conditions. Our studies implicate the importance of 6S RNA in stabilizing RNAP interactions with σ^A^, and suggest that it exerts its main roles in prolonged stationary phase. Our findings give insight into the mode of action of 6S RNA in an important pathogen, and suggest the need to develop strategies that prevent low level rifampicin from persisting in the antibiotic-treated host.

This protective effect is possibly due to steric hindrance, as the presence of 6S RNA would reduce the accessibility of the RNAP to its inhibitors. A second non-exclusive proposal is that the destabilization of σ^A^ associated with the absence of 6S RNA affects the transcriptional program to adapt to low concentrations of RNAP inhibitors. Note that in *S. aureus*, this shielding effect is not associated with the sigma stress factor σ^B^.

## Material and methods

### Bacterial strains and culture

All strains used in this study and their genotypes are listed in Table S1. Strains were cultured at 37°C, with 180 rpm agitation for liquid cultures except for *C. difficile.* The latter was cultured in anaerobic conditions (5% H_2_, 5% CO_2_, 90% N_2_) with 7.5 μg.mL^-1^ (precultures) or 15 μg.mL^-1^ thiamphenicol (plates) for plasmid selection. *E. coli* and *S. enterica* (serovar Typhimurium) strains were cultured in lysogeny broth (LB), *S. aureus* strains in brain heart infusion (BHI) or tryptic soy broth (TSB) and *C. difficile* in BHI. When necessary, media were supplemented with antibiotics.

*S. aureus* mutants were constructed in HG003 or HG002 background (59) by allelic exchange using pIMAY (77) derivatives (except strains for RNAP purification) as described (33). Plasmids used in this study are described in Table S2. Most plasmids constructed for this study were obtained by Gibson assembly (78) as described using primers from Table S3, and cloned in *E.coli* IM08B (79). Genetic features of the main *S. aureus* mutants used in this study are represented in Fig.S1.

To purify *S. aureus* σ factors, *sigA* and *sigB* from HG003 were PCR amplified using primers F-SigA/R-SigA-His and F-SigB/R-SigB-His respectively, and cloned into the NdeI/XhoI restriction sites of pET-21C vector. The resulting pET-21C-sigA and pET-21C-sigB plasmids were transformed into *E. coli* strain BL21(DE3) pLysS leading to strains producing σ^A^ and σ^B^ (His)_6_-tagged in their C-terminal portions upon IPTG induction.

A HG003 strain expressing a chromosomally encoded His-tagged RpoC for the purification of the RNAP core enzyme was constructed as followed. The recombinational transfer of the Histidine sequence into *rpoC* gene was achieved by two-steps PCR. A sequence encoding for ten Histidines was added upstream from the termination codon, in frame with RpoC (β’ subunit of RNA polymerase). The *rpoC-His* fragment was generated by long-flanking homology PCR using the primers listed in table S3 and cloned between the BamH1 and PstI restriction sites of temperature-sensitive pBT2 vector (80) to obtain pBT2-rpoC-His plasmid. The resulting plasmid was electroporated into *S.aureus* RN4220 and then transferred to HG003 strain. The gene encoding for His-tagged RpoC protein was integrated in the *S. aureus* HG003 chromosome by double cross-over recombination as described (81) to obtain the HG003 *rpoC-his* strain.

*S. enterica* serovar Typhimurium mutants were constructed in the background of strain LT2-derived MA7455 (82) using λ Red recombineering (83). See Fig.S2 for constructions.

*C. difficile* mutants were constructed in 630Δ*erm* background (84). The knock out mutant was obtained using an allelic chromosomic exchange following the published method (85) with primers CM57/CM58 and CM59/CM60 and pMSR vector pDIA7052. To complement, *ssrS^Cd^* sequence and its promoter region were PCR amplified using the primer pair CM77/CM78 and cloned into pMTL84121 to produce pDIA7065. Two *E. coli* strains were used as intermediate: NEB 10-Beta for plasmid construction (Table S2) and HB101 RP4 for conjugation.

### MIC determination

Antimicrobial susceptibility testing by broth microdilution for rifampicin, rifabutin, fidaxomicin and aureothricin MIC determination in *S. aureus* were performed as described (86). *S*. *enterica* serovar Typhimurium rifampicin MIC was determined as described (87).

### Fitness experiment

The experiment was performed as described (33) with three independent sRNA tagged-mutant libraries grown simultaneously in TSB +/- 6 μg.L^-1^ rifampicin and sampled at four points: OD_600_=1, O/N, OD_600_=1 after the first dilution and OD_600_=1 after the second dilution. All the mutants were tag-sequenced with an adapted Illumina protocol. The amount of each mutant was normalized to the total amount of bacteria with/without rifampicin and to the inoculum (Fig. 1A). Totally, three mutants (*locus1, rsaD(tag26*) and *teg146*) were discarded from the analysis because of experimental troubleshooting. Concerning *sprD* and *sau5949*, only two values were taken in account in the 3^rd^ dilution sampling.

### Spot test

Overnight cultures were 10-fold serial diluted in NaCl 0.9% (*S. aureus* and *S*. enterica) or BHI (*C. difficile)* until 10^-8^ and spotted on agar plates containing different sublethal antibiotic concentrations, namely less than MIC (rifampicin < 12 μg.L^-1^ for *S. aureus* or < 12 μg.mL^-1^ for *S.* enterica, rifabutin < 15.6 μg.L^-1^, aureothricin < 6.25 μg.mL^-1^, fidaxomicin < 4 mg.L^-1^ for *S. aureus* or < 30 μg.L^-1^ for *C. difficile*). Pictures were taken after O/N growth or 24h for *C. difficile*.

### Growth curves

*S. aureus* strains were cultured in microplates from O/N triplicates cultures 1/1000 diluted in BHI +/- 5μg.L^-1^ rifampicin. Two and three day-cultures were also used as pre-cultures in Fig. 6C. Absorbance at 600 nm (OD_600_) was measured overtime with a plate reader (Clariostar).

### Fluorescence measurement

mAmetrine expression (λ_exc_ = 425 +/- 15 nm, λ_em_ = 525 +/- 15 nm) was monitored overtime in microplates by a plate reader (Clariostar), simultaneously to absorbance measurement of overnight triplicates cultures diluted 1:1000 in TSB to limit auto-fluorescence.

### RNA extraction

Strains were cultured until the desired OD_600_. After centrifugation, pellets were frozen in dry-ice ethanol. RNAs were then extracted by phenol-chloroform treatment as described (88). When necessary, RNAs were incubated with Turbo DNase treatment (Thermo Fisher Scientific) prior to a second phenol-chloroform extraction.

### Northern Blot

10 μg total RNAs per well were separated on 1.3% agarose or 10% TBE-urea polyacrylamide gel (Criterion Precast gel) as described (89). For polyacrylamide gels, electrophoresis in TBE 1X was followed by transfer to Hybond-N^+^ membrane in TBE 0.5X using TE70 ECL Semi-Dry Transfer Unit (Amersham Pharmacia Biotech). Probes (Table S3) were [α-^32^P]dCTP-labelled.

### RNA-seq and transcriptomic analysis

RNAs (DNA-free) from triplicate cultures sampled at OD_600_= 7 were sequenced using NextSeq 500/550 High Output Kit v2 (75 cycles). Sequences were aligned to the reference genome (CP000253-1-NCTC8325) with Bowtie2 tool and quantified with Feature Counts program. Differential gene expression analysis was performed using DESeq2 algorithm (90). The data for this study have been deposited in the European Nucleotide Archive (ENA) at EMBL-EBI under accession number PRJEB50160 (https://www.ebi.ac.uk/ena/browser/view/PRJEB50160).

### 5’-3’ RACE

5’ and 3’ ends were determined using circularization method described in (51). 10 μg total RNA of wild-type strain (HG003) were extracted from samplings at OD_600_ 7, O/N and day 4 (D4). Primers used to amplify the 5’/3’ junction with Phusion high-fidelity DNA polymerase (Thermo Fisher Scientific) are listed in Table S3. CloneJET PCR cloning kit (Thermo Fisher Scientific) was used to clone the final PCR products.

### Purification of σ^A^, σ^B^ and RNAP core enzyme

For σ purification, *E. coli* strains BL21(DE3) pET-21C-*sigA* and BL21(DE3) pET-21C-*sigB* were grown in 1 L of LB broth at 37°C to OD_600_ 0.5. After induction with 1 mM isopropyl-1-thio-β-d-galactopyranoside for 3 h, bacteria were collected by centrifugation, resuspended in 10 mL of buffer A (10 mM HEPES, pH 7.5, 200 mm NaCl, 1mM MgCl_2_, 20 mM Imidazol, 5% glycerol) with 0.25 mg.mL^-1^ lysozyme per gram of pellet and frozen and thawed two times. Lysates were treated with DNase I (100 units.mL^-1^) for 20 min at 30°C, and supernatants containing σ^A^-His or σ^B^-His proteins were obtained by centrifugation at 8000 g for 15 min. For RNA polymerase purification, a fresh overnight culture of *S. aureus* HG003 *rpoC-his* was used to inoculate 500 mL of BHI at OD_600_ 0.1 and grown 5 hours at 37°C. The culture was harvested by centrifugation at 4000 g for 15 min. The cell-pellet was resuspended in 10 mL of buffer A with 1 mg of lysostaphin and DNase I (100 units.mL^-1^). After an incubation of 20 min at 37°C, the lysate was clarified by centrifuging 30 min at 40000 g. For affinity purification of σ^A^-His, σ^B^-His and (His)10-tagged RNAP, HisTRAP^TM^ TALON column (5 mL, GE healthcare) was connected to an AKTA prime chromatography system (GE healthcare) equilibrated with buffer B (10 mM HEPES, pH 7.5, 1 M NaCl, 20 mM Imidazol, 5 % glycerol). After loading the lysate containing either σ^A^-His, σ^B^-His or (His)_10_-tagged RNAP, the column was washed with 100 mL of buffer B. His-tagged proteins were then eluted using an imidazole gradient, dialyzed in buffer C (10 mM HEPES, pH 7.5, 100 mM NaCl, 1 mM MgCl_2_, 5 % glycerol) and then subjected to a second step of purification on heparin column. Proteins were loaded on a HiTrap Heparin HP column (1 mL, GE healthcare) equilibrated with buffer C. After a wash with 20 mL of buffer C, proteins were eluted using a NaCl gradient, dialyzed in buffer C and then concentrated in a centrifugal concentrator with a 10-kDa molecular weight cutoff membrane (Merck Millipore).

### Electrophoretic mobility shift assay (EMSA)

*ssrS^Sa^* and *sprB* were *in vitro* transcribed from a PCR product template T7 promoter-containing (primers listed in Table S3) with MEGAscript T7 transcription kit (Thermo Fisher Scientific). RNAs were separated on 8 % polyacrylamide-7 M urea gel electrophoresis and eluted overnight in G50 elution buffer (20 mM Tris-HCl pH7.5, 2 mM EDTA and 0.25 % SDS). RNAs were precipitated in cold ethanol and 0.3 M of sodium acetate and dephosphorylated using Calf-intestinal alkaline phosphatase (New England Biolabs), according to manufacturer protocol. Obtained RNAs were 5’-radiolabelled with T4 polynucleotide kinase (New England Biolabs) and [γ^32^P] adenosine triphosphate (ATP) and purified with MicroSpin G-50 column (Amersham Pharmacia Biotech). 140 nM of RNAP alone or preincubated 10 minutes at 37°C with 420 nM of Sigma factors were mixed with 4 nM of radiolabelled 6S RNA or SprB in buffer D (15 mM Hepes pH 7.5, 100 mM NaCl, 1 mM MgCl_2_, 5 % glycerol, 100 μg.mL^-1^ BSA, 200 μg.mL^-1^ *E. coli* tRNA). Complex formation was performed at 37°C during 10 min and samples were loaded on 5 % polyacrylamide-5 % glycerol gel under non-denaturing conditions. Gels were dried and visualized using a Typhoon Phosphorimager (Molecular Dynamic).

### σ^A^ and β subunits quantification

*S. aureus* strains were cultured in triplicates until the desired OD_600_. Frozen pellets were lysed in 50 mM TRIS-HCl buffer pH 7.5 with glass beads. Total protein amount in the supernatant was determined by Bradford protein assay. Western-Blot electrophoresis was performed with 3 μg proteins per well, using 8% Bis-TRIS Plus polyacrylamide gel (Bolt, Invitrogen). Transfer and hybridization followed iBlot and iBind manufacturer instructions (Invitrogen) respectively. Membranes were pre-hybridized at 4°C O/N with human serum 1/10000e to saturate unspecific binding. Rabbit primary antibodies were used for immunodetection of σ^A^ (anti-σ^A^, 1:5000 dilution) and β subunit (anti-RNAP, 1:10000 dilution). A HRP-conjugated goat anti-rabbit (Advansta, 1:4000 dilution) was chosen as secondary antibody. Pictures were taken with a CCD Camera.

## Acknowledgments

We are grateful to our colleague Sandy Gruss (INRAE, MICALIS) for critical reading of the manuscript. We thank Patricia Kerboriou (I2BC), Claire Toffano-Nioche (I2BC) and Mehdi El Sadek Fadel (I2BC) for technical, bioinformatics and statistical support. We are grateful to Masaya Fujita (University of Houston) who graciously provided anti-σ^A^ antibodies. We thank Dodo Bourbon for helpful discussions and warm support. We acknowledge the high-throughput sequencing facility of I2BC for its sequencing and bioinformatics expertise and for its contribution to this study.

This work was supported by the Agence Nationale de la Recherche [ANR-19-CE12-0006-01 (RRARE)]. M.E. and W. L were the recipient of scholarships from the *Ministère de l’Enseignement supérieur, de la Recherche et de l’Innovation* (MESRI) and Chinese scholarship council (CSC), respectively.

